# Comparable neutralization of SARS-CoV-2 Delta AY.1 and Delta in individuals sera vaccinated with BBV152

**DOI:** 10.1101/2021.07.30.454511

**Authors:** Pragya D. Yadav, Rima R Sahay, Gajanan Sapkal, Dimpal Nyayanit, Anita M. Shete, Gururaj Deshpande, Deepak Y. Patil, Nivedita Gupta, Sanjay Kumar, Priya Abraham, Samiran Panda, Balram Bhargava

## Abstract

The recent emergence of the SARS-CoV-2 Variant of Concern, B.1.617.2 (Delta) variant and its high transmissibility has led to the second wave in India. BBV152, a whole-virion inactivated SARS-CoV-2 vaccine used for mass immunization in India, showed a 65.2% protection against the Delta variant in a double-blind, randomized, multicentre, phase 3 clinical trial. Subsequently, Delta has been further mutated to Delta AY.1, AY.2, and AY.3. Of these, AY.1 variant was first detected in India in April 2021 and subsequently from twenty other countries as well. Here, we have evaluated the IgG antibody titer and neutralizing potential of sera of COVID-19 naive individual’s full doses of BBV152 vaccine, COVID-19 recovered cases with full dose vaccines and breakthrough cases post-immunization BBV152 vaccines against Delta, Delta AY.1 and B.1.617.3. A reduction in neutralizing activity was observed with the COVID-19 naive individuals full vaccinated (1.3, 1.5, 1.9-fold), COVID-19 recovered cases with full BBV152 immunization (2.5, 3.5, 3.8-fold) and breakthrough cases post-immunization (1.9, 2.8, 3.5-fold) against Delta, Delta AY.1 and B.1.617.3 respectively compared to B.1 variant. A minor reduction was observed in the neutralizing antibody titer in COVID-19 recovered cases full BBV152 vaccinated and post immunized infected cases compared to COVID-19 naive vaccinated individuals. However, with the observed high titers, the sera of individuals belonging to all the aforementioned groups they would still neutralize the Delta, Delta AY.1 and B.1.617.3 variants effectively.

## Text

The severe acute respiratory syndrome coronavirus-2 (SARS-CoV-2) Delta variant (B.1.617.2), a variant of concern (VOC), was associated with the second wave in India during March-May 2021^1^. The Delta variant outpaced the other two sub-lineages Kappa (B.1.617.1) and B.1.617.3 and was responsible for 90% of the cases reported in India^2^. It has spread across nearly in 99 countries and found to be more infectious than the Alpha, Beta and Gamma variants. The recent reports suggested that irrespective of reduced neutralization, the Delta variant was found to be susceptible to BNT162b2, BBV152/Covaxin and Covishield vaccines^3–5^. However, the Delta variant has also been identified as the leading cause of breakthrough infections globally among vaccinated individuals^6^.

Delta variant with its characteristic spike protein mutations (L452R, T478K, D614G and P681R) has mutated further into four sub-lineages with an additional mutations which are associated with higher transmission and probable immune escape^1^. Recently, Delta variant has mutated to Delta AY.1, AY.2 and AY.3. Of these, apparently highly infectious Delta AY.1 variant was first detected in India in April 2021 and subsequently from twenty other countries as well^1^. Cases of Delta AY.1 have mostly been reported from Europe, Asia and America with the highest prevalence observed in Portugal, Japan, United States of America, United Kingdom and Switzerland^7^. The variant has characteristic mutations in the genome at ORF1a (A1306S, P2046L, P2287S, V2930L, T3255I, T3646A), ORF1b (P314L, G662S, P1000L, A1918V), S (T19R, L452R, T478K, D614G, P681R), ORF3a (S26L), M (I82T), ORF7a (V82A, T120I), ORF7b (T40I), ORF8 (del119/120) and N (D63G, R203M, G215C, D377Y) to the increasing speculation about its ability to escape immune response, has been a major concern for the ongoing vaccination programs^8^. Moreover, this variant contains an additional K417N mutation in the spike protein and emerging evidence suggest that this mutation could lead to resistance against monoclonal antibodies i.e., Casirivimab and Imdevimab^9^. However, it is uncertain whether Delta AY.1 is capable of causing higher transmissibility, severe disease and evasion of immune response compared to the Delta variant. Till now, the prevalence of Delta AY.1 is found to be low in India and the rest of the world and, no information is available on the efficacy of currently available vaccines against the Delta AY.1 variant.

The clinical efficacy against COVID-19 infection of BBV152, a whole-virion inactivated SARS-CoV-2 vaccine was assessed in a double-blind, randomized, multicentre, phase 3 clinical trial on 25,798 participants to evaluate the efficacy, safety, and immunological lot consistency of BBV152. Efficacy against asymptomatic COVID-19 was 63.6% and 65.2% protection against the SARS-CoV-2 Variant of Concern, B.1.617.2 (Delta)^10^.

Here, we present the neutralization activity of the sera of individuals vaccinated with two doses of BBV152 [n=42; Female (n=24; median age:43.5yrs); Male (n=18; median age:46 yrs)] collected between 2.5 to 22 weeks [median: 16 weeks] after second dose, COVID-19 recovered cases with two doses of BBV152 [n=14; female (n=8; median age: 44.5yrs); male (n=6; median age: 42 yrs)]14-70 weeks [median: 38 weeks] after second dose and breakthrough cases post two-dose BBV152 vaccinations [n=30; female (n=17; median age: 45 yrs); male (n=13; median age: 39 yrs)] collected between 2-18 weeks (median: 9 weeks) against Delta, Delta AY.1, and B.1.617.3, compared to B.1 variant. Neutralizing antibody (NAb) titers of all the serum specimens against all the variants were determined using a 50% plaque reduction neutralization test^11^. Besides this, IgG response was also assessed using whole inactivated SARS-CoV-2 antigen, nucleocapsid and S1-RBD protein IgG ELISA.

The sera of individuals with two dose vaccination showed a geometric mean titer (GMT) of NAb to be 310.6 (95% confidence interval (CI): 222-434.6); 241.6 (95% CI: 167.8-347.7); 209.1 (95% CI: 146.5-298.3); 165.3 (95% CI: 115.6-236.5) against B.1, Delta, Delta AY.1 and respectively. The sera of the recovered cases with two dose of vaccine showed rise in NAb titer against B.1 had GMT 820.1 (95% CI: 469-1434), Delta 328.6 (95% CI: 186.9-577.9), Delta AY.1 234.5 (95% CI: 138.7-396.4) and B.1.617.3 217.8 (95% CI: 136.7-347.1). Sera from breakthrough cases had higher NAb titer compared to the individuals from the other two groups mentioned before. The GMT titers were 896.6 (95% CI: 550.3-1461), 465.6 (95% CI: 213.2-1016), 317.2 (95% CI: 125.5-801.4), 259.7 (95% CI: 157.1-429.4) against B.1, Delta, Delta AY.1 and B.1.617.3 respectively (Figure 1a-d).

**Figure 1:**
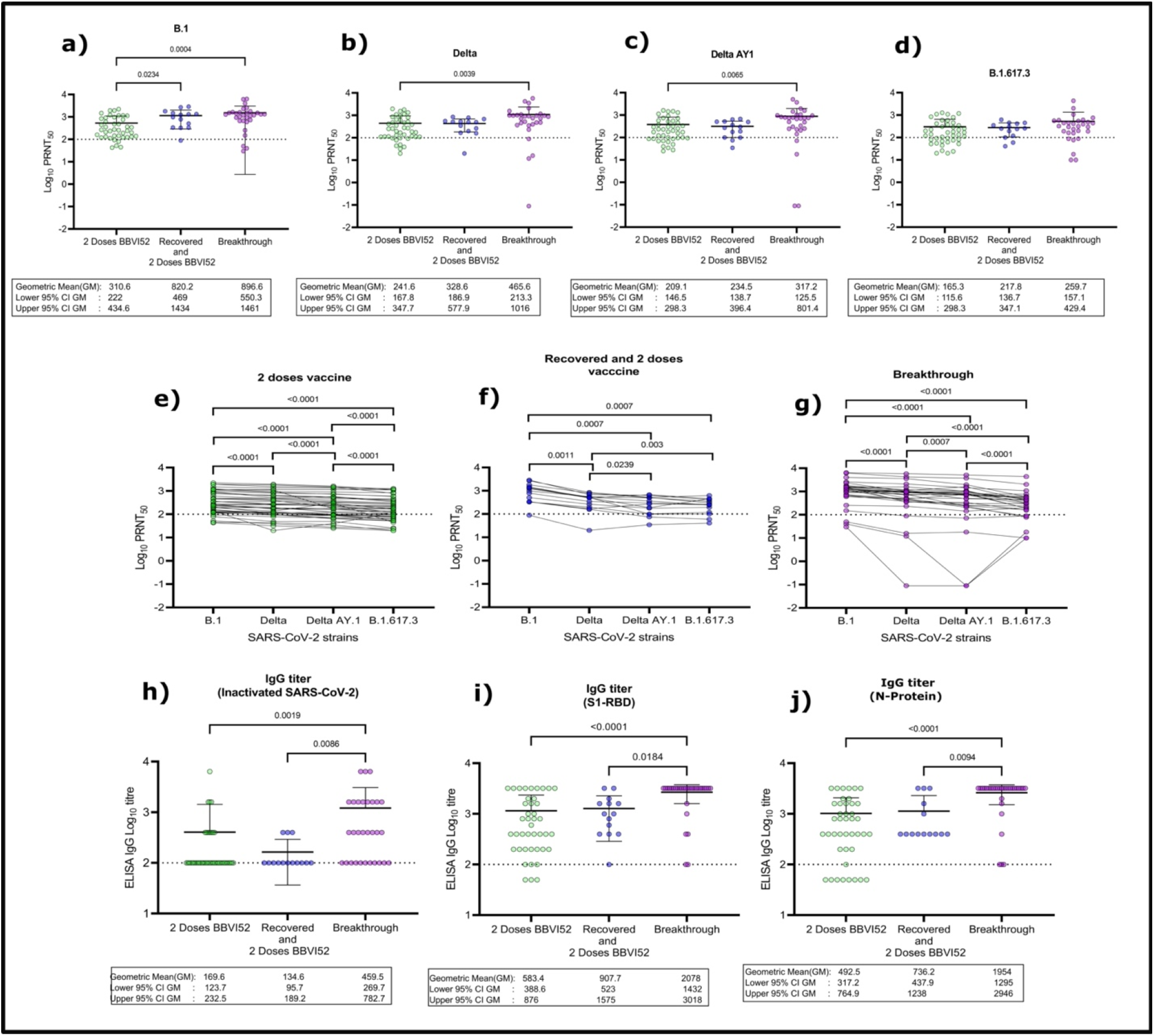
Neutralization of individual sera vaccinated with BBV152 vaccines from different scenarios against B.1, Delta, Delta AY.1 and B.1.617.3 strains and ELISA titer of individual sera vaccinated from different scenarios. Neutralization titer comparison of individual cases sera immunized with two dose vaccine BBV152, recovered case sera immunized with two dose vaccine BBV152 and breakthrough cases; B.1(GISAID identifier: EPL_ISL_825088) (a), delta (GISAID accession number: EPI_ISL_2400521) (b), delta AY1 (GISAID accession number EPI_ISL_2671901) (c), and B.1.617.3 (GISAID accession number: EPL_ISL_2497905) (d). The statistical significance was assessed using a two-tailed Kruskal Wallis test with Dunn’s test of multiple comparisons was performed to analyze the statistical significance. NAb titer of individual sera vaccinated with two doses of BBV152 (e), recovered cases administered with two doses of BBV152 (f) and breakthrough cases (g).A matched pair two-tailed pair-wise comparison was performed using the Wilcoxon signed-rank test to analyze the statistical significance. Anti-SARS-CoV-2 IgG titers of vaccinated individual’s sera for inactivated SARS-CoV-2 (h), S1-RBD protein (i) and N protein (j). The statistical significance was assessed using a two-tailed Kruskal-Wallis test with Dunn’s test of multiple comparisons. P-value less than 0.05 were considered to be statistically significant for the tests applied. The dotted line on the figures indicates the limit of detection of the assay. Data are presented as mean values +/− standard deviation (SD).

The sera of individuals who were fully immunized (with 2 doses) didn’t show significant fold-reduction in the NAb titer against Delta, Delta AY.1 and B.1.617.3 [1.29 (p-value: < 0.0001); 1.49 (p-value : <0.0001); 1.88 (p-value: <0.0001)] compared to B.1 (Figure 1e). However, the sera of recovered cases with vaccinations and breakthrough cases had considerable fold-reductions in the NAb titer of [2.5 (p-value: 0.0011); 3.5 (p-value: 0.0007); 3.77 (p-value: 0.0007)] and [1.93 (p-value: <0.0001); 2.83 (p-value: <0.0001); 3.45 (p-value: <0.0001)] against Delta, Delta AY.1 and B.1.617.3 variants respectively compared to the prototype strain (Figure 1f-g). However, NAb titer of sera of recovered cases with vaccinations and breakthrough cases were significantly high compared to two dose vaccinations.

The anti-SARS-CoV-2 ELISA of the vaccinated individual’s sera samples from the breakthrough individual’s had 2.7 fold (p-value: 0.0019), 3.56 fold (p-value: <0.0001) and 3.97 fold (p-value: <0.0001) higher than the other individual sera receiving 2 doses of vaccine for inactivated SARS-CoV-2, S1-RBD protein and N protein respectively (Figure 1h-j).

A significant increase in the NAb titer against B.1 variant in recovered cases with vaccination and breakthrough cases was observed compared to the Covid-19 naive vaccinees. Similarly, a significant increase in NAb titer was also observed among these two groups against Delta, Delta AY.1 and B.1.617.3 variants. This demonstrates the possible role of memory cells in immune boosting with post-infection or infection after immunization. The comparative analysis of all the groups revealed that the B.1.617.3 variant seems to be less susceptible to neutralization followed by Delta AY.1 and Delta variants compared to B.1 (Figure 1a-g). A recent study demonstrated a reduction in neutralization by 4 fold and 11 fold against Delta variant with the sera of healthy individuals vaccinated with two doses of ChAdOx1 and BNT162b2 vaccine respectively^8^. Our earlier studies with BBV152/Covaxin™ and Covishield™ have shown 2.7- and 3.2-fold reduction in NAb titer against Delta variant compared to B.1^4,5^. The present study revealed 1.5, 3.5, 2.8-fold reduction in NAb titer for the Delta AY.1 Sera of vaccinees Covid naïve, recovered cases with full vaccination and breakthrough cases demonstrated 1.3, 2.5 and 1.9-fold reduction against Delta variant in comparison to B.1 variant respectively.

In conclusion, reduction in the NAb titer was observed in the sera of fully immunized Covid naïve, recovered cases with BBV152 vaccination and breakthrough cases. However, with the observed high titers, the sera of individuals belonging to all the aforementioned groups they would still neutralize the Delta, Delta AY.1 and B.1.617.3 variants effectively.

## Supporting information

Supplementary Information Methodology

## Ethical approval

The study was approved by the Institutional Human Ethics Committee of ICMR-NIV, Pune, India under the project ‘Assessment of immunological responses in breakthrough cases of SARS-CoV-2 in post-COVID-19 vaccinated group’.

## Author Contributions

PDY contributed to study design, data analysis, interpretation and writing. RRS, GS, GRD, DAN, AMS, DYP and SKY contributed to data collection, data analysis, interpretation and writing. PA, NG, SP, and BB contributed to the critical review and finalization of the paper.

## Acknowledgement

We sincerely acknowledge the excellent support of Mr. Prasad Sarkale, Mr. Shreekant Baradkar, Dr. Rajlaxmi Jain, Ms. Aasha Salunkhe, Mr. Chetan Patil, Mrs. Triparna Majumdar and Mrs. Savita Patil during the study.

