## Supplementary Information Methodology for "Comparable neutralization of SARS-CoV-2 Delta AY.1 and Delta in individuals sera vaccinated with BBV152"

**Anti-SARS-CoV-2 IgG antibody evaluation**

We evaluated anti-SARS-CoV-2 IgG antibody response against S1-RBD, N-protein and the whole antigen. Briefly, 96-well ELISA plates (Nunc, Germany) were coated with SARS-CoV-2 specific antigens (S1-RBD at a concentration of 1.5µg/well and N protein at a concentration of 0.5µg/well in PBS pH 7.4). The plates were blocked with a Liquid Plate Sealer (CANDOR Bioscience GmbH, Germany) and Stabilcoat (Surmodics) for two hours at 37°C. The plates were washed twice with 10 mM PBS, pH 7.4 with 0.1 per cent Tween-20 (PBST) (Sigma-Aldrich, USA). The sera of were serially diluted four-fold and added to antigen-coated plates and incubated at 37°C for one hour. These wells were washed five times using 1× PBST and followed by addition 50 μl/well of anti-human IgG horseradish peroxidase (HRP) (Sigma) diluted in Stabilzyme Noble (Surmodics). The plates were incubated for half an hour at 37°C and then washed as described above. Further, 100 μl of TMB substrate was added and incubated for 10 min. The reaction was stopped by 1 N H2SO4, and the absorbance values were measured at 450 nm using an ELISA reader. Anti-SARS-CoV-2 antibody (NIBSC code 20/130) was also included in the assay as a reference standard. The cut-off for the assays was set at twice of average OD value of negative control. The endpoint titer of a sample was defined as the reciprocal of the highest dilution that had reading above the cutoff value.

**Neutralizing antibody evaluation**

The plaque reduction neutralization assay (PRNT50) was performed against the B.1, Delta (B.1.617.2), Delta AY.1 and B.1.617.3 strains. The sera were serially diluted four-fold and mixed with an equal amount of B.1, Delta (B.1.617.2), Delta AY.1 and B.1.617.3 virus suspension (50-60 Plaque forming units in 0.1 ml) separately. Post incubation at 37°C for 1 hr, each virus-diluted serum sample (0.1 ml) was inoculated onto duplicate wells of a 24-well tissue culture plate of Vero CCL-81 cells. After incubating the plate at 37°C for 1 hr, an overlay medium consisting of 2% Carboxymethyl cellulose (CMC) with 2% fetal calf serum (FCS) in 2× MEM was added to the cell monolayer. The plate was further incubated at 37°C with 5% CO2 for 5 days. At assay termination, plates were stained with 1% amido black for an hour. NAb titers were defined as the highest serum dilution that resulted in >50 (PRNT50) reduction in the number of plaques.
